# Synthesis of Glycoconjugates Utilizing the Regioselectivity of a Lytic Polysaccharide Monooxygenase

**DOI:** 10.1101/2020.03.13.990838

**Authors:** Bjørge Westereng, Stjepan K. Kračun, Shaun Leivers, Magnus Ø. Arntzen, Finn L. Aachmann, Vincent G. H. Eijsink

## Abstract

Polysaccharides from plant biomass are the most abundant renewable chemicals on Earth and can potentially be converted to a wide variety of useful glycoconjugates. While anomeric hydroxyl groups of carbohydrates are amenable to a variety of useful chemical modifications, selective cross-coupling to non-reducing ends has remained challenging. Several lytic polysaccharide monooxygenases (LPMOs), powerful enzymes known for their application in cellulose degradation, specifically oxidize non-reducing ends, introducing carbonyl groups that can be utilized for chemical coupling. This study provides a simple and highly specific approach to produce oxime-based glycoconjugates from LPMO-functionalized oligosaccharides. The products are evaluated by HPLC, mass spectrometry and NMR. Furthermore, we demonstrate potential biodegradability of these glycoconjugates using selective enzymes.

## Introduction

Over the years, numerous methods for synthesis of glycoconjugates have been developed, mainly using traditional synthetic methods. An unprotected carbohydrate contains multiple hydroxyl groups with similar reactivity. Coupling ligands to a specific position is therefore challenging and normally requires an exhaustive number of chemicals and synthetic steps^1,2,3^. This is one of the major hurdles for efficient utilization of some of Nature’s most abundant molecules as building blocks for novel biomolecules. Glycoconjugates have a considerable potential as drugs, fine chemicals and in material science^4^. Moreover, the efficiency of enzymatic depolymerization of plant polysaccharides has improved tremendously in recent years, partly due to the inclusion of oxidative enzymes in enzyme mixtures^5,6,7^. This development enables more efficient production of plant derived carbohydrate-based “bulk chemicals”, including a wide variety of structurally diverse oligosaccharides that could be converted to glycoconjugates.

Oxidative carbohydrate-active enzymes are involved in a multitude of biological processes, including biomass decay. These enzymes use different mechanisms to generate oxidations on mono-, oligo- and/or polysaccharides. These oxidation reactions produce either carbonyls or carboxylic acids and, for poly- and oligomeric substrates, these reactions involve enzymes such as LPMOs^5,8,9,10^, cellobiose dehydrogenase (CDH)^11,12^, galactose oxidase^13^, pyranose dehydrogenase^14,15^ and glucooligosaccharide oxidases^16^. Together, these enzymes enable a range of oxidative functionalizations^17^ and offer several possibilities for subsequent glycoconjugation^18^. Notably, there are very few enzymes that oxidize the non-anomeric hydroxyls in the sugar rings of oligo- and polysaccharides.

LPMOs may play a special, albeit so far scarcely explored, role in glyco-functionalization, since they are active on polymers and can functionalize polymeric surfaces. The initial step of the lytic action of LPMOs on cellulose entails breaking the energetically strong C-H bond^19^ at position C1 or C4, catalyzed by a triangularly shaped catalytic copper site termed the “histidine brace”^9^. Some LPMOs oxidize C1, thus generating a carboxylic acid that could also be generated in alternative ways, for example using CDH. Interestingly, other LPMOs have the ability to specifically oxidize C4, generating a ketone functionality that is hard to generate in other ways. Due to the discovery of an increasing number of C4-oxidizing LPMOs active on abundant polysaccharides such as cellulose^20^, glucomannan^21^, xylan^22^ and xyloglucan^21^, several target substrates for non-reducing end functionalization by LPMOs are now available.

A feasible way to synthesize heteropolymers is via polycondensation (analogous to the synthesis of well-known materials like polyesters and polyurethanes). This requires carbohydrates that are bi-functionalized. While functionalization and/or cross coupling to carbohydrate reducing ends is relatively straightforward, doing the same at the non-reducing end has proven more challenging. The possibility to specifically functionalize C4 using LPMOs opens a possible route towards generating bi-functional carbohydrate building blocks.

Here we describe a highly selective chemo-enzymatic coupling procedure for the C4 position of oligosaccharides derived from one of the most abundant polysaccharides in Nature, cellulose (Scheme 1 sketches the different synthetic steps). We further demonstrate that this allows the production of enzymatically cleavable bi-functional glycoconjugates.

## Results and discussion

Several LPMOs characterized so far have mixed activities leading to a combination of C1- and C4-oxidized products, whereas some strict C4-oxidizers act on oligomeric substrates and thus produce short products^23^. These properties would interfere with the coupling chemistry and complicate analysis in this proof of concept study. Hence, we searched for an LPMO that is not active on oligosaccharides and only releases C4-oxidized products with a degree of polymerization (DP) that would provide oligosaccharides of a suitable range (3-6). Based on previous studies, the C4-oxidizing *Neurospora crassa* enzyme, *Nc*LPMO9A (NCU02240)^24,25^, which is not active on soluble cellooligosaccharides, was selected. More detailed analysis, necessary to validate this enzyme’s suitability for the present study, showed that it releases longer, oxidized oligosaccharides (predominantly DP 3-5) (Fig. 1). Furthermore, MS/MS analysis displayed in Fig. S1 showed fragmentation patterns typical of C4-oxidized cello-oligosaccharides^23^.

**Figure 1.**
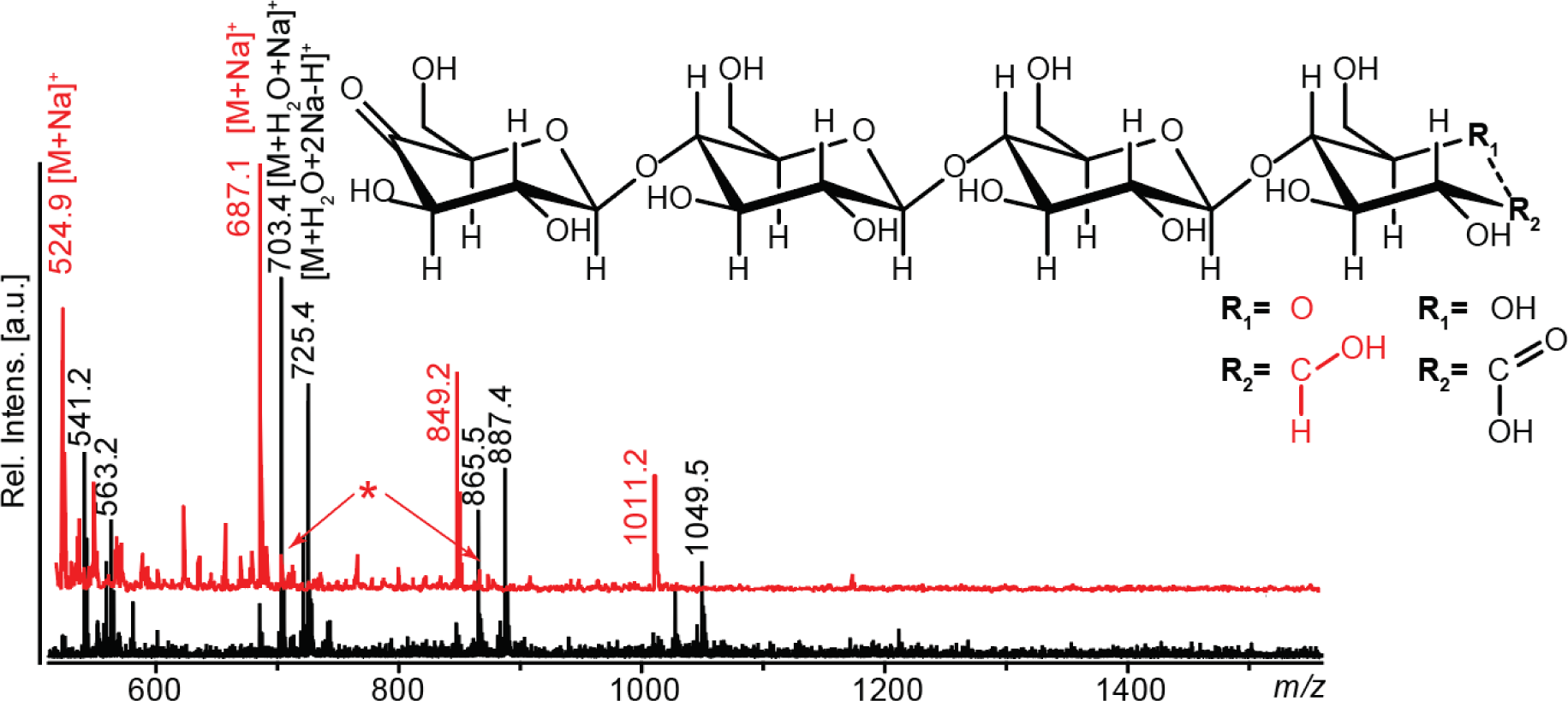
MALDI ToF MS analysis of products generated from PASC by *Nc*LPMO9A using ascorbic acid (red structure and spectrum) or CDH as reductant (black). The reaction with CDH will lead to oxidation of reducing ends and thus generate doule oxidized products. The main peaks in the spectra correspond to (*m/z* values apply to sodium adducts of the tetramer): 687, mono-oxidized (keto-sugar; dominates in the red spectrum); 703, double oxidized (aldonic acid form at C1; dominates in the black spectrum); 725, double oxidized (sodium salt of the aldonic acid form; dominates in the black spectrum). The following minor species are also visible: 685, double oxidized (keto at C4, lactone form at C1); 705, single oxidized (gemdiol of 4-keto sugar, indicated by arrows and *). Similar peak clusters were observed for other oligomers.

When *Nc*LPMO9A was combined with a cellobiose dehydrogenase (*Mt*CDH from *Myrococcum thermophilum*^37^, the dominating ketone signal in the MALDI-ToF MS spectra shifted by 16 or 38, representing the formation of an aldonic acid (+16), which subsequently has a tendency to form a double sodium adduct (M-H+2Na; +38) (Fig. 1). Since CDH oxidizes oligosaccharides in the reducing end, this observation confirms that *Nc*LPMO9A oxidizes in the non-reducing end only.

Since it is challenging to determine the oxidative regioselectivity of LPMOs by HPAEC and MALDI^26,27^, we designed a precise and simple way for probing C4-oxidation on oligosaccharides, inspired by previous work by Beeson et al.^20^ who looked at monosaccharides. Reduction of the ketone at C4 results in a mixture of galactose and glucose and hence reduction of a C-4 oxidized cello-oligosaccharide will result in the formation of a mixture of oligosaccharides with either a glucosyl or a galactosyl at the non-reducing end and a glucitol (GlcOH) in the reducing end. To assess this in detail, we first generated GalGlc_n_ (n=3 or 4) standards (Fig. 2) by using UDP-Gal (donor), cellotriose/tetraose (acceptor) and a galactosyltransferase. These oligosaccharides have considerably shorter elution times in HPAEC than their corresponding cello-oligosaccharides (Fig. 3A). Oligosaccharides generated by *Nc*LPMO9A and our in-house prepared oligomeric standards were reduced to completion using NaBD_4_ and the resulting oligomeric products were analyzed directly by HPAEC and MALDI ToF MS (Figs. 2, 3B and 3C). Reaction products generated by *Nc*LPMO9A indeed yielded a mixture of (Glc)_n_GlcOH and Gal(Glc)_n_GlcOH (n=2 to 4) oligosaccharides confirming C4 oxidation. These results further prove that *Nc*LPMO9A only oxidizes C4.

**Figure 2.**
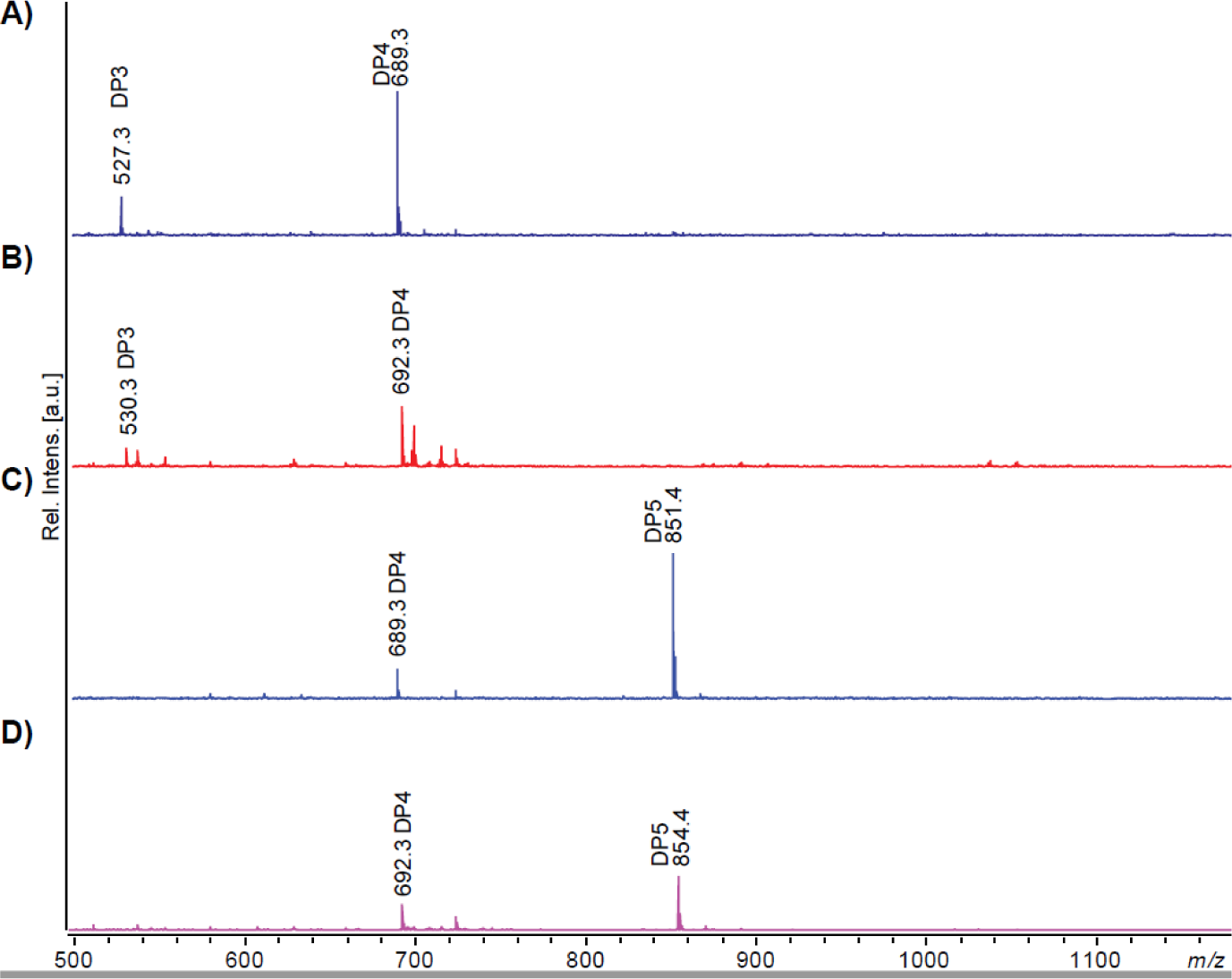
MALDI ToF MS of GalGlc_n_ standards, prior to and after reduction with NaBD_4_. The non-reduced compounds shown in (A), GalGlc_3_ (at *m/z* 689 with a little bit of unreacted Glc_3_ at *m/z* 527), and (C), GalGlc_4_ (at *m/z* 851 with a little bit of unreacted Glc_4_ at *m/z* 689), yield masses equivalent to cello-oligosaccharides. The reduced compounds (B & D) show an expected *m/z* of +3, reflecting reduction and the incorporation of one D (see Fig. 3 for structures).

**Figure 3.**
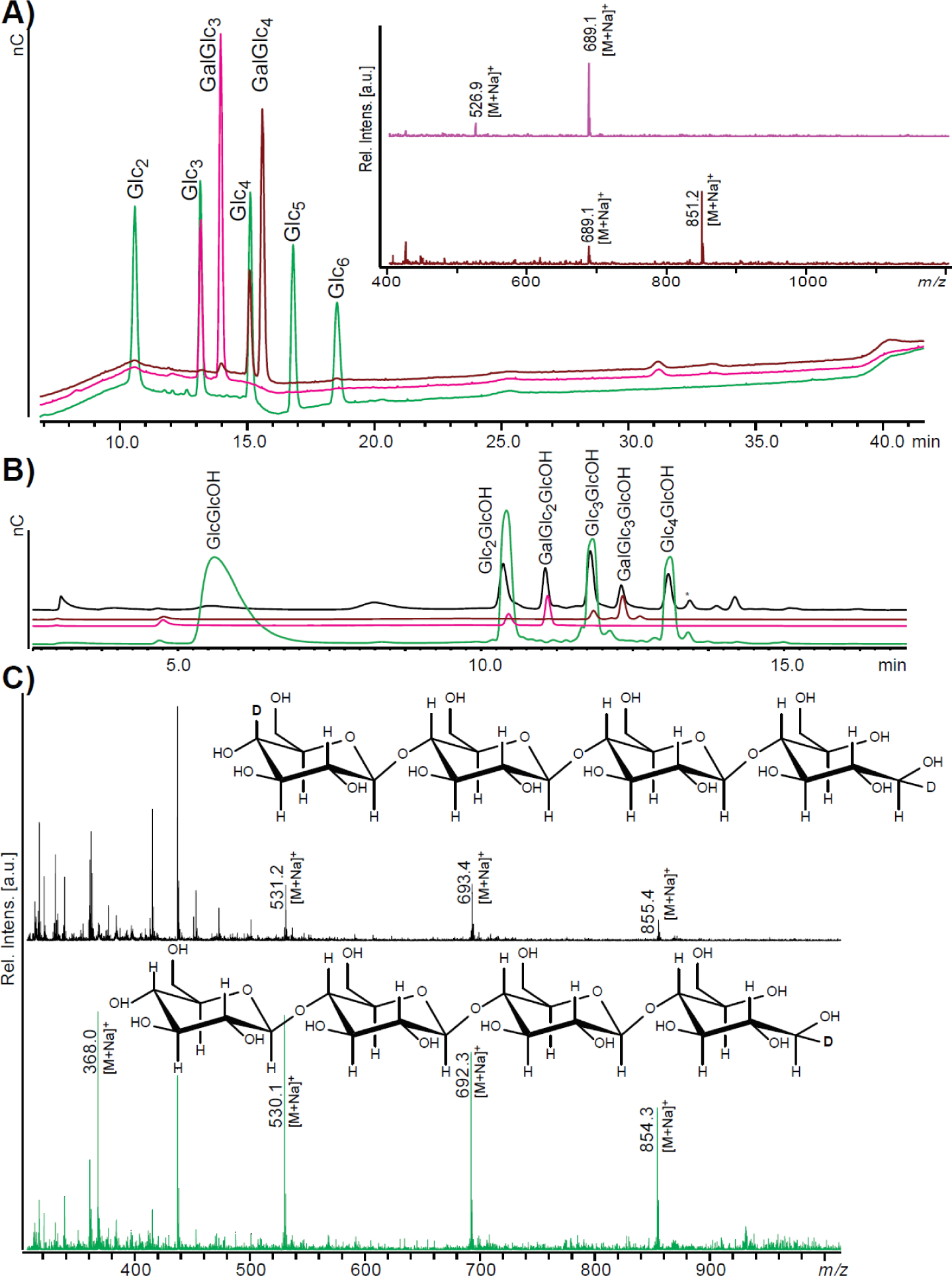
Reduction of soluble products generated by *Nc*LPMO9A. A) Superposition of HPAEC chromatograms of various standards, showing that GalGlc_3_ and GalGlc_4_ have weaker retention than cello-oligosaccharides of the same DP. The inset to the right shows the MALDI ToF-MS spectrum for GalGlc_3_ (pink, *m/z* 689 with a little bit of unreacted Glc_3_ at *m/z* 527) and GalGlc_4_ (brown, *m/z* 851 with a little bit of unreacted Glc_4_ at *m/z* 689). B) HPAEC chromatograms of oligosaccharides reduced with NaBD_4_: Glc_2-5_ (green), GalGlc_3_ (pink), GalGlc_4_ (brown) and products generated by *Nc*LPMO9A (black). Abbreviations: GlcOH, glucitol, Glc, glucose and Gal, galactose C) MALDI ToF MS of products generated by *Nc*LPMO9A (black) and a DP2-5 mixture of cello-oligosaccharides (green) each reduced by NaBD_4_. In the C4-oxidized products generated by *Nc*LPMO9A one extra D is incorporated; expected *m/z* values for the tetrameric compound are: native, 689; C4-oxidized, 687; native reduced with NaBD_4_, 689 + 2 + 1 = 692; C4-oxidized reduced with NaBD_4_, 687 + 2 + 1 + 2 + 1 = 693; Insets show the structures of the DP4 compounds.

Notably, this reduction approach, used here to proof C4 oxidation, also provides a relatively simple method for quantification of oxidized products, based on using NaBD_4_ for reduction. After reduction, oligosaccharides with oxidized C4 will possess two deuteriums whereas non-oxidized oligosaccharides will only contain one deuterium, thus yielding an easily detectable difference of *m/z* = 1 (Fig. 3C). Internal standards could thus be produced from non-oxidized oligosaccharides as described above, possibly using NaBH_4_ for reduction, generating a more pronounced difference of *m/z* = 2. Fig. 1 and 3 further show that *Nc*LPMO9A primarily releases oligosaccharides of DP3-5, which is a suitable size for the purpose of this study. Several experiments were performed in order to evaluate the specificity of *Nc*LPMO9A. Screening of other substrates using MALDI-ToF MS for product detection showed that, in contrast to several other C4 oxidizing LPMO9s, *Nc*LPMO9A is not active on curdlan [β-(1,3) glucan], pustulan [β-(1,6) glucan], β-chitin, glucomannan, xylan, xyloglucan, mixed linkage glucan [β-(1,3),(1,4)] and is thus highly specific for cellulose. Furthermore, there was no activity on cello-oligosaccharides with a degree of polymerization (DP) of 2 to 6, which shows that this enzyme is not active on solubilized oligomeric products. Together the properties demonstrated above makes *Nc*LPMO9A suitable for this proof of concept study on C4-functionalized cellooligosaccharides.

Oxidized products generated by *Nc*LPMO9A (C4-oxidized) or by a combination of *Nc*LPMO9A and CDH (C1 & C4-oxidized; Fig. 1) were subjected to targeted carbonyl functionalization using a hetero-bi-functional linker (aminooxy-linker), 2- (aminooxy)-1-ethanaminium dichloride, to generate oximes (Fig. 4). Product formation was confirmed by MALDI-ToF MS (Fig. 4) and porous graphitized carbon electrospray ionization mass spectrometry (PGC-ESI-MS) (Fig. 5) as well as by NMR (Fig. 6). Due to the presence of a carbonyl function in the reducing end, oxime formation takes place at both ends of the C4-oxidized oligosaccharide. Chromatography/tandem mass spectrometry (LC-MS/MS) with PGC yielded masses corresponding to three double-oxime functionalized oligosaccharides (*m/z* 619.25, 781.28 and 943.35). Furthermore, the obtained fragmentation patterns could be matched with the oxime functionalized oligosaccharides (Fig. 5). Notably, when analyzing the oxime functionality on PGC we observed a considerably increased retention of the oligosaccharides (Fig. 5), compared to the corresponding C-4 oxidized oligosaccharides^27^. Replacing ammonium acetate with a stronger eluent like formic acid in the gradient was needed to enable elution.

**Figure 4.**
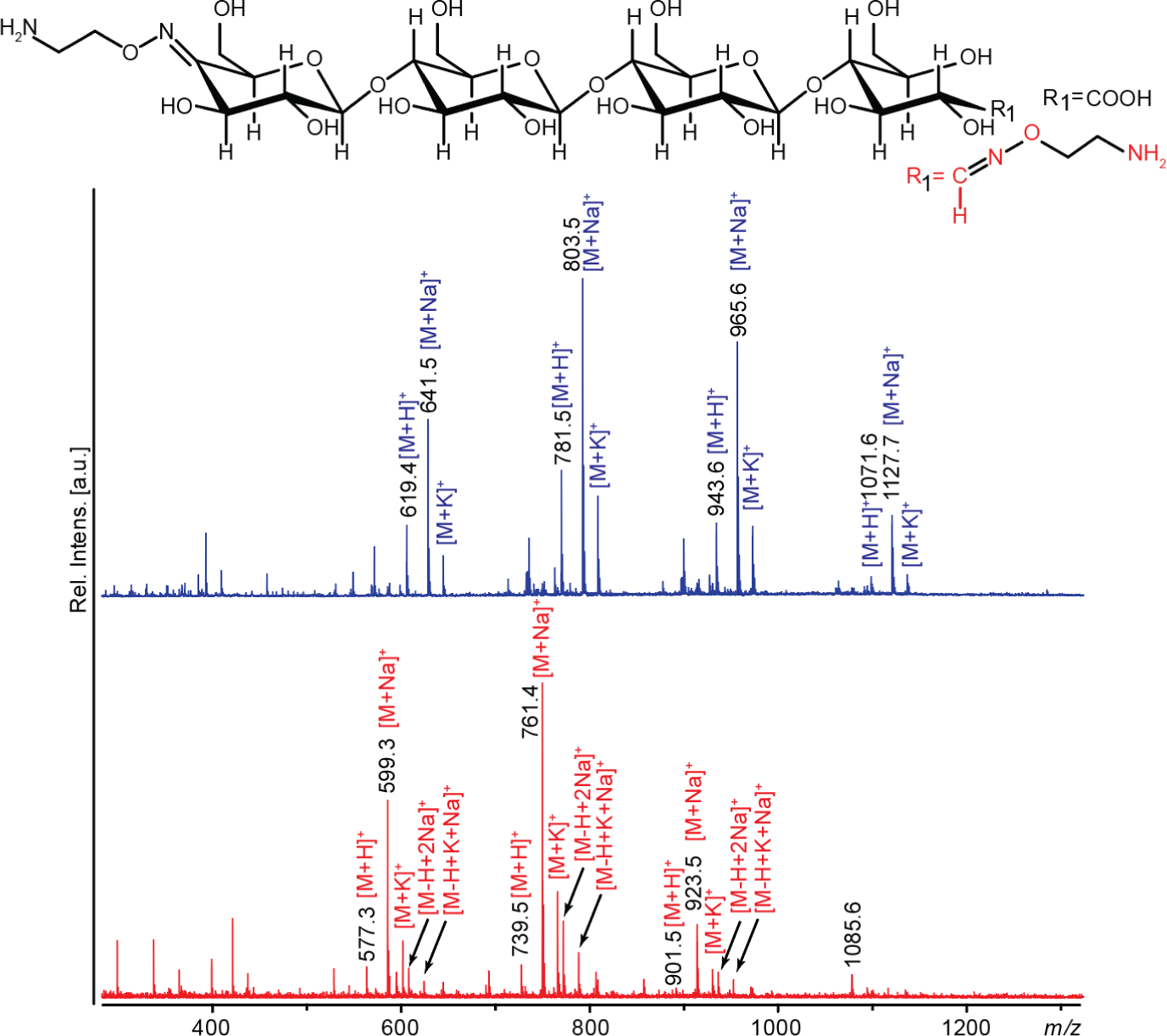
MALDI ToF MS spectra of oxime functionalized oligosaccharides. The structure of a functionalized tetramer is shown above the spectra. The upper spectrum shows bi-functionalized oligosaccharides obtained from a product mixture generated by *Nc*LPMO9A from PASC. The spectrum shows clear signals for a trimeric, tetrameric, and pentameric double-funtionalized compound and a lower signal for the hexameric product (*m/z* 641, 803, 965 and 1127, respectively; all sodium adducts). There are also peaks corresponding to proton and potassium adducts denoted [M+H]^+^ and [M+K]^+^. The lower panel shows functionalization of LPMO-generated C4-oxidized products that were C1-protected through oxidation by CDH (*m/z* values are 42 lower compared to panel A; sodium adducts). The latter spectrum also shows peaks corresponding to [M-H+2Na]^+^ and [M-H+K+Na]^+^ adducts, which are formed due to the presence of an (CDH-generated) aldonic acid group.

**Figure 5.**
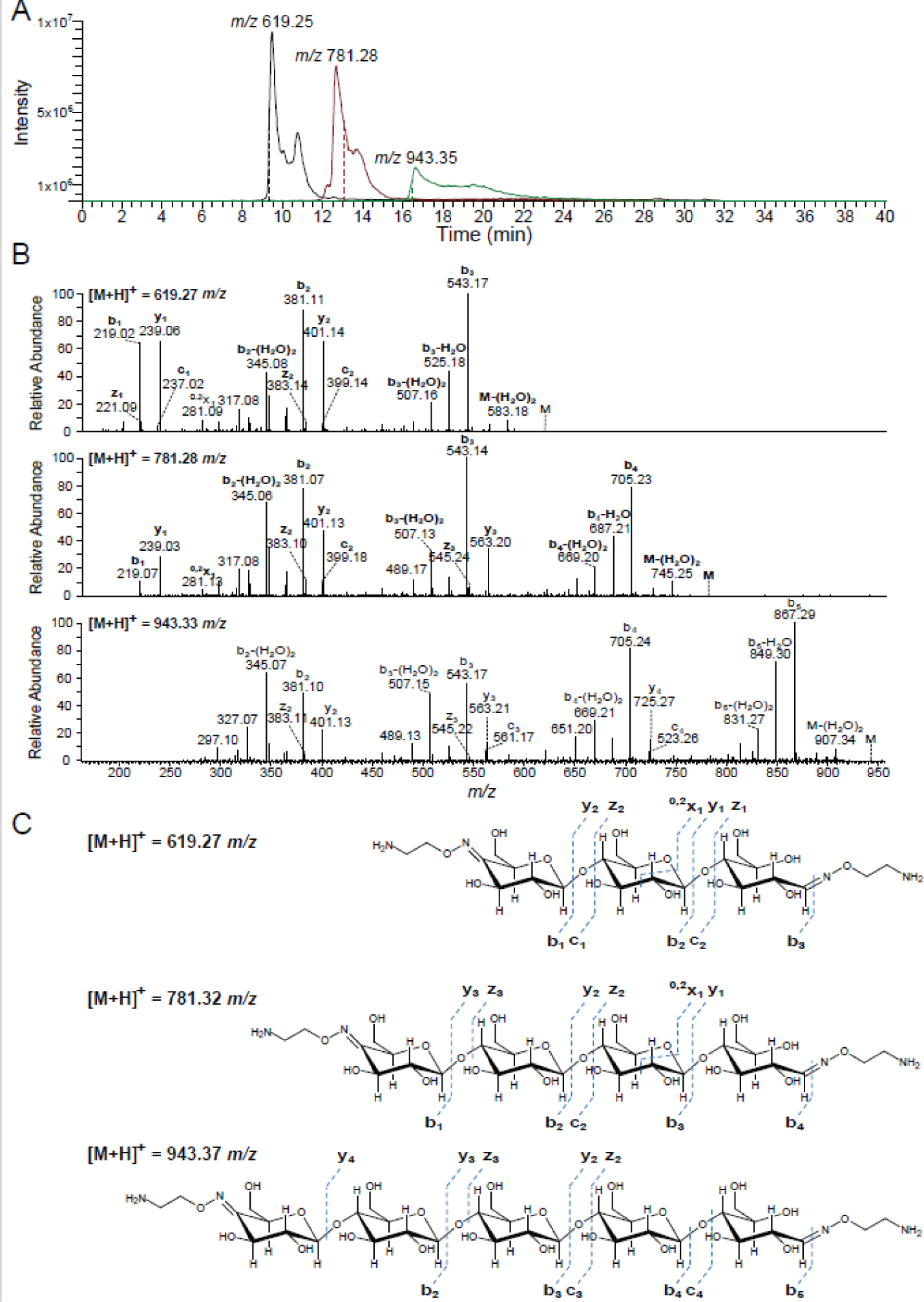
PGC-MS analysis of bi-functionalized cello-oligosaccharides. The oxime bi-functionalized oligosaccharides were separated by porous graphitized carbon (PGC) chromatography using an LTQ Velos Pro mass spectrometer (Thermo) for detection. A) Overlay of three extracted ion chromatograms for *m/z*-ratios 619.25, 781.28 and 943.35, showing separation of the bi-functionalized products with DP 3, 4 and 5, respectively (note that these are proton adducts). The dashed line shows the time of recording of the MS/MS spectra. B) MS/MS spectra of the three products show a dominant cleavage of the glycosidic bonds followed by multiple water losses (annotated as B-(H2O)2). A cleavage of the reducing-end amine linker bond can also be seen resulting in the following fragments; 543.17, 705.23 and 867.29 for the mother ions 619.25, 781.28 and 943.35, respectively. C) Structures of the three bi-functionalized products, with theoretical masses and the observed fragmentation patterns (using the nomenclature of Domon and Costello)^46^.

**Figure 6.**
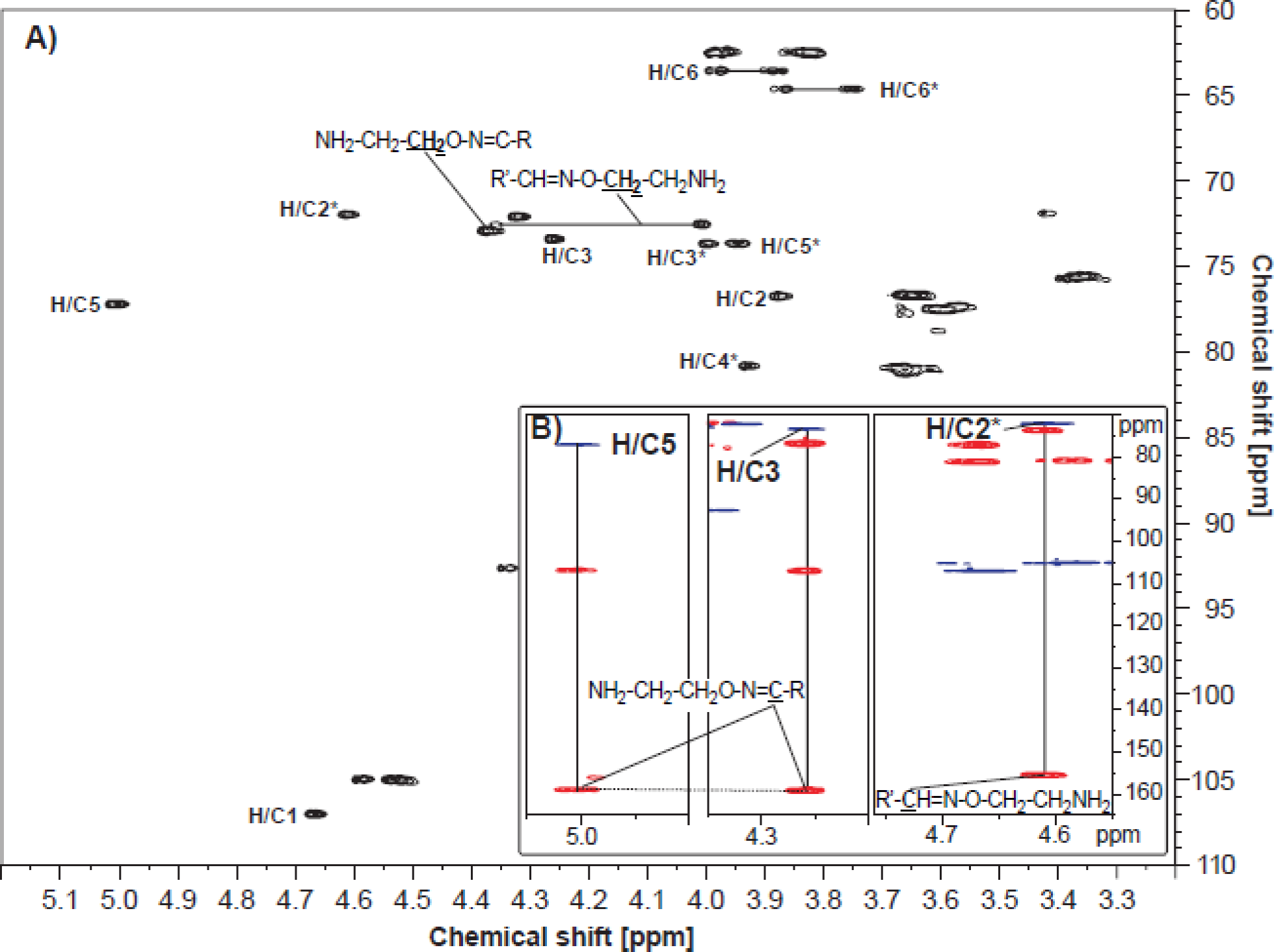
NMR analysis of bi-functional cello-oligosaccharides. A) [^1^H-^13^C] HSQC spectrum; the sample was in 99.9% D_2_O and the spectrum was recorded at 25 °C. Peaks in the proton/carbon signals of the C4ox residue in the non-reducing end are marked by H/C#, where # indicates the carbon number in the residue. Peaks in the proton/carbon signals of the Glc1 (= the former reducing end) spin system are marked by H/C#*. Chemical groups giving rise to signals for the 2-aminooxy group are in bold and underlined. For the sake of simplicity, peaks related to internal monosaccharide residues are not marked (a full assignment of the chemical shifts is provided in Table S1). B) Correlations between a [^1^H-^13^C] HSQC spectrum and a [^1^H-^13^C] HMBC spectrum recorded for the bi-functionalized product. The left and the middle insert show correlations (indicated by vertical lines) from H/C-3 and H/C-5 peaks in the HSQC (blue) for the C4-oxidized end to peaks (indicated by a horizontal dotted line) with a common carbon chemical shift of 159.2 ppm in the HMBC (red). The right insert shows a correlation from H/C-2* in the HSQC (blue) for the reducing end to a carbon peak with a carbon chemical shift of 155.4 ppm in the HMBC (red). Both carbon chemical shifts obtained from the HMBC spectrum correspond well to the (expected) presence of an imine (-C=N-) group, which is expected to be formed during the coupling reaction (Fig. 5), thus confirming the structure of the bi-functionalized product. R and R’ represent the non-reducing and the reducing end of the bi-functionalized cello-oligosaccharides, respectively. Further confirmation of the successful coupling can be found in the DOSY spectrum, Figure S2.

In double oxidized oligosaccharides, containing a CDH-generated carboxyl functionality that will block oxime formation at the former “reducing end”, only the carbonyl function at C4 in the non-reducing end will react. Indeed, for C4-oxidized oligosaccharides that were protected in their reducing end by CDH-catalyzed oxidation to an aldonic acid, only one oxime was formed, in the non-reducing end. The efficiency of the reaction was assessed by mass spectrometry (Figs 4 and 5) and the position of the oxime was verified by NMR both for the reducing and the non-reducing end (Fig. 6, Fig S2, Table S1).

Structural elucidation by NMR provided direct proof of the structure of the putatively bi-functionalized products generated by coupling aminooxy-linkers to either one or both ends oligosaccharides. The individual monosaccharides were assigned by starting at the anomeric signal as well as from the primary alcohol group at C6 and then following ^1^H-^1^H connectivity using DQF-COSY, H2BC and [^1^H-^13^C] HSQC-[^1^H,^1^H] TOCSY (full assignment of shift values in Table S1). Most of the carbon chemical shifts were obtained from [^1^H-^13^C] HSQC (Fig. 6). The [^1^H-^13^C] HMBC spectrum provided long range bond correlations that connect the monosaccharides to the aminooxy-linker. The carbon chemical shift of the C4 oxidized end was determined by combining the correlations from a [^1^H-^13^C] HSQC spectrum and a [^1^H-^13^C] HMBC spectrum (see Fig. 6B). Both for the reducing end (Glc1 – H/C-2) and for the C4 oxidized non-reducing end (C4ox-3 and C4ox-5) HMBC correlations are observed from proton signals 4.61 ppm and 5.01/4.27 ppm to a carbon signal at 155.4 ppm and 159.2 ppm, respectively (see Fig. 6B). These carbon chemical shifts fit well to the expected carbon chemical shifts for an aminooxy group, which would form upon coupling. Altogether, the NMR results suggest that the structure of the bi-functional products correspond with the structures depicted in Fig. 5C.

The aminooxy-functionalized oligosaccharides can be enzymatically degraded using a cellulase, to regenerate a native reducing and non-reducing end, while producing shorter oligosaccharides, one or both of which may be oxime-functionalized, depending on the starting compound (Fig. 7). This effectively expands the application potential of the present technology and underpins the flexibility of these conjugates. Two different cellulases were used to demonstrate degradability of the functionalized oligosaccharides (Fig. 7). Both Cel6B from *T. fusca* and Cel5A from *H. jecorina* degraded the aminooxy-functionalized oligosaccharides. Both enzymes degraded bi-functionalized DP5 and DP6 completely, and Cel5A also degrade most of bi-functionalized DP4. Since the mass difference of degradation products with the oxime linker in the reducing end is *m/z* +2 compared to products where the oxime linker occurs in the non-reducing end, the mass spectra reveal preferred sites for cellulase cleavage. For example, Cel5A primarily converted the double-functionalized pentamer to a trimer with C4-oxime linker (583.2) and a dimer with a reducing end linker (423.1), whereas conversion to a dimer with C4-linker (421.1) and a trimer with a reducing end linker (585.2) was less frequent. For Cel6B the degradation pattern for the double-functionalized pentamer was slightly different; here the main products were a trimer with a reducing end linker (585.2) and a dimer with a C4-linker (421.1). This exemplifies a rational approach for generating defined functionalized oligosaccharides. Essentially, by playing with the initial reaction (one or two oxime functionalizations) and the use of cellulases, in principle any combination of a natural and a modified chain end can be generated. More uniform products may be obtained by first purifying the C4-oxidized cello-oligosaccharides, which could be carried out using Porous Graphitized Carbon columns or HILIC ^27^.

**Figure 7.**
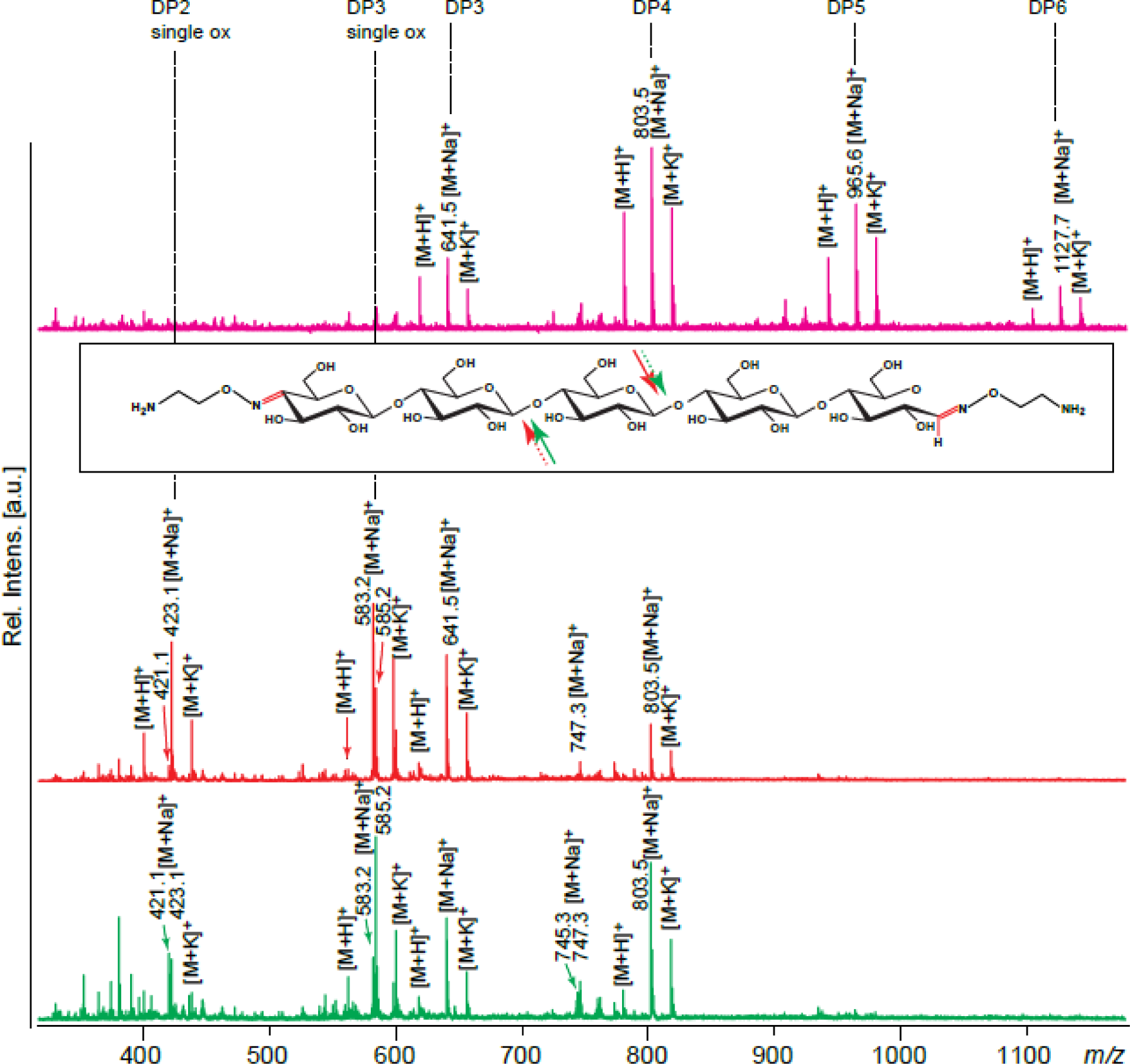
MALDI-ToF-MS analysis of a mixture of bi-functionalized aminooxy-cello-oligosaccharides and products formed after cellulase treatment of this mixture. The spectra shown are for an untreated mixture (magenta), a Cel5A treated mixture (red spectrum and red arrows; enzyme reaction 3) and a Cel6B treated mixture (green spectrum and green arrows; enzyme reaction 4). The insert shows the structure of the bi-functionalized pentamer, with solid and dotted arrows indicating the major and minor cleavage activity of Cel5A (red) and Cel6B (green), respectively; see text for details. Adducts of proton, sodium and potassium are labelled as [M+H]^+^, [M+Na]^+^ and [M+K]^+^, respectively.

By combining C4-specific LPMOs and CDHs, more complex glycoconjugates may be generated since one may employ chemistries pertaining to carbonyl groups, such as the method described above, and chemistries pertaining to carboxyl groups, for example using carbodiimide activation^28^. Notably, C1-specific LPMOs, or even LPMOs with a mixed C1/C4 activity could also be considered for this purpose. Importantly, the use of LPMOs in principle allows partial surface oxidation of polysaccharide fibers^29, 30, 49^, which may enable functionalization of fiber surfaces without affecting the tensile strength of the fibers. Such applications could be of commercial interest.

The functional group introduced through aminooxy functionalization was an amino group, and the usefulness of this group for further functionalization of the glycoconjugate was demonstrated by labeling with fluorescein isothiocyanate (FITC) that is highly reactive towards amines. Indeed, after FITC labeling, oligosaccharides with oxime functionalization in both ends carried either one (due to incomplete labeling) or two FITC molecules (Fig. 8A, Fig. 9 B-E). On the other hand, labeling oligosaccharides oxidized by CDH and thus carrying only one oxime functionalization carried only a single FITC specifically labeled at C4 in the non-reducing end (Fig. 8B, Scheme 1). The aminooxy functionalization shown in this study demonstrated only one of many possibilities for functionalization. Other functional groups such as azides and alkynes are also feasible and will extend the potential of the approach described here.

**Figure 8.**
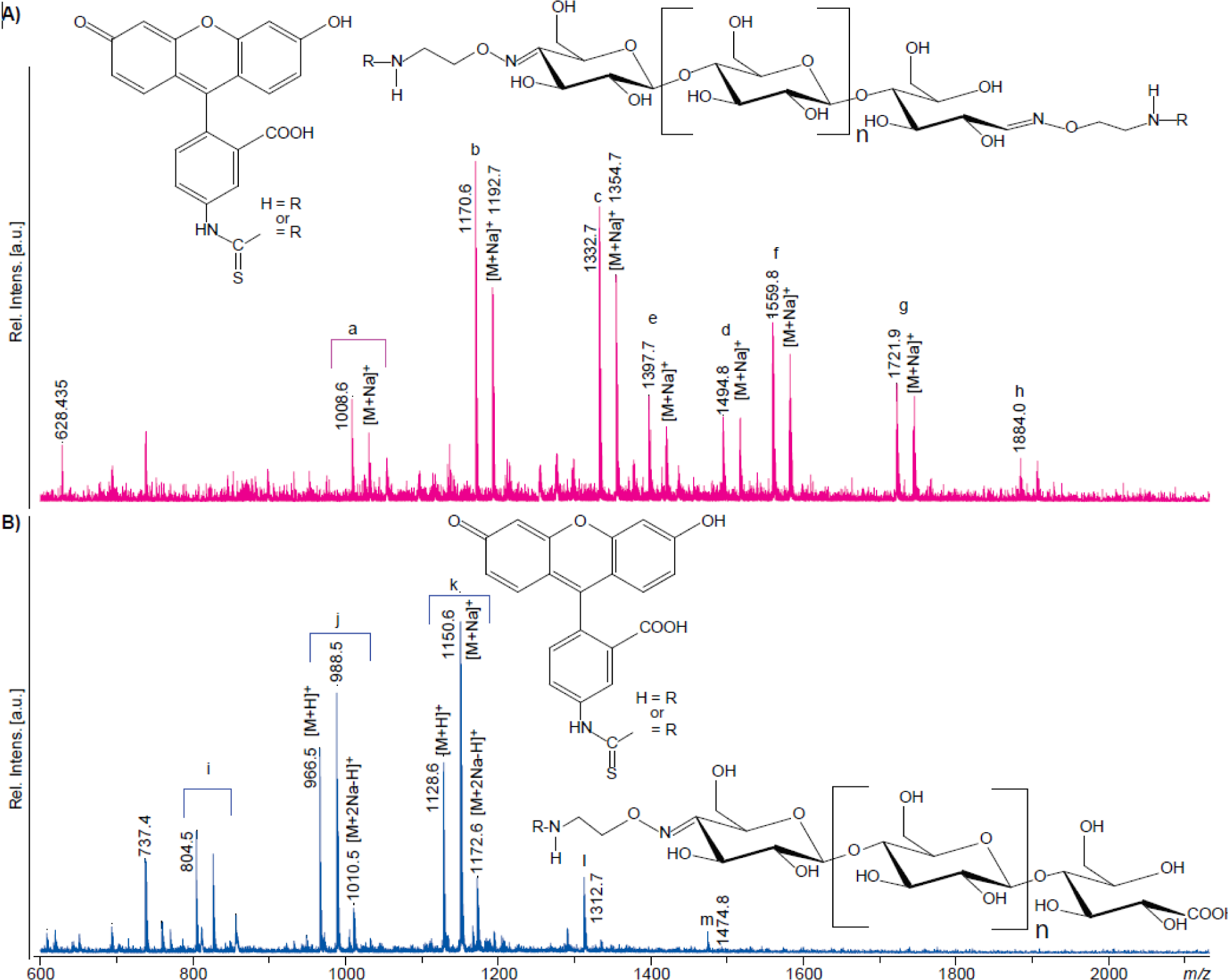
MALDI-ToF-MS analysis of the results of FITC labeling. A) FITC labeling of double oxime-functionalized oligosaccharides with possible structures shown above the mass spectrum. Peak clusters representing single labeled products are a (n=1), b (n=2), c (n=3) and d (n=4), and peak clusters representing double labeled products are e (n=1), f (n=2), g (n=3), and h (n=4). This figure shows that labeling with FITC was incomplete, meaning that also singly-labeled products were present. This is most likely due to the use of a short reaction time (1h) and a low amount of FITC equivalents. B) FITC labeling of single oxime-functionalized oligosaccharides with possible structures shown to the lower right. The signal clusters represent i (n=0), j (n=1), k (n=2), l (n=3) and m (n=4). Note the double sodium which is typical for saccharides containing carboxylic acids^47,48^. Adducts of proton, sodium and sodium salts of sodium adducts are labelled as [M+H]^+^, [M+Na]^+^ and [M+2Na-H]^+^, respectively. R corresponds to FITC or H.

**Figure 9.**
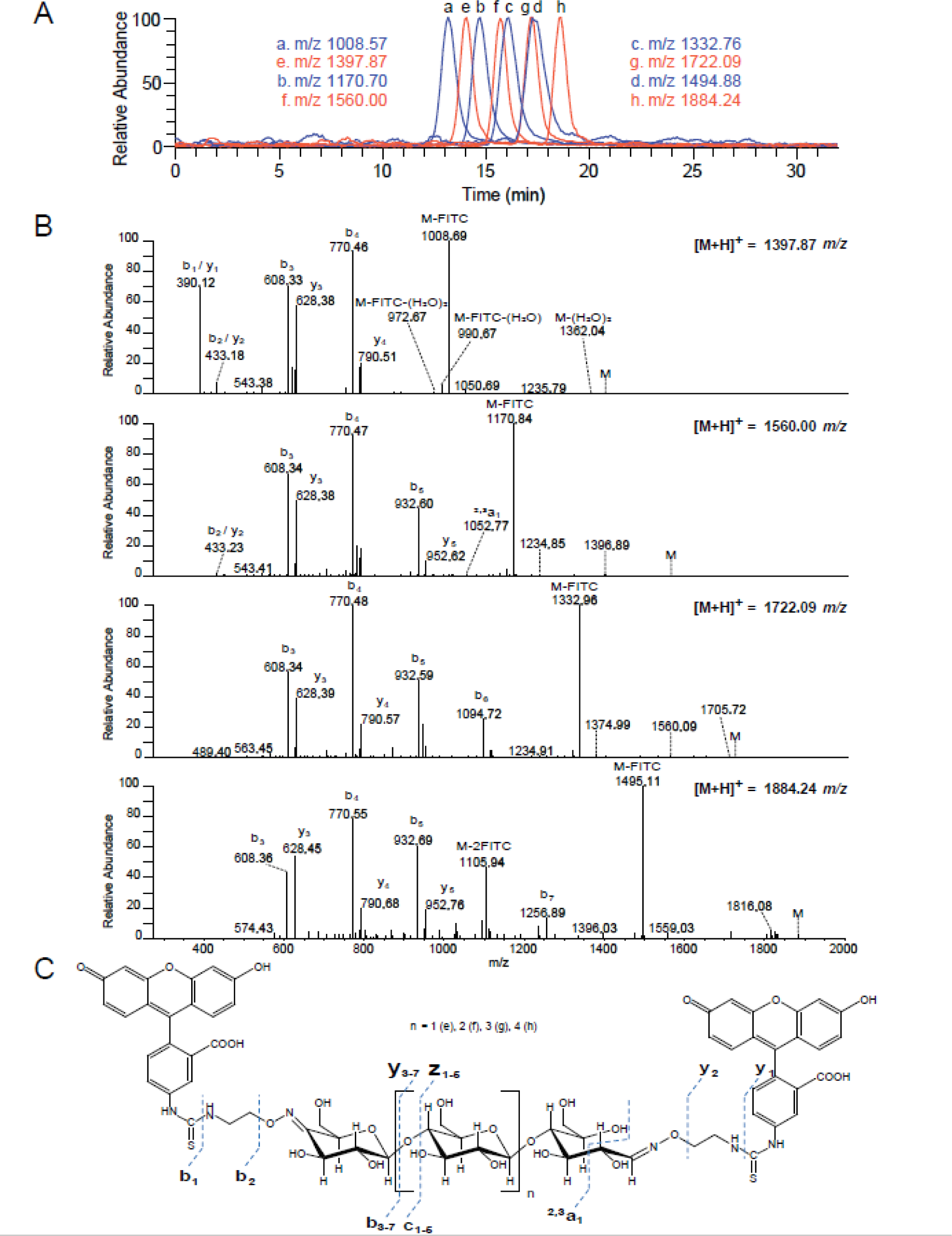
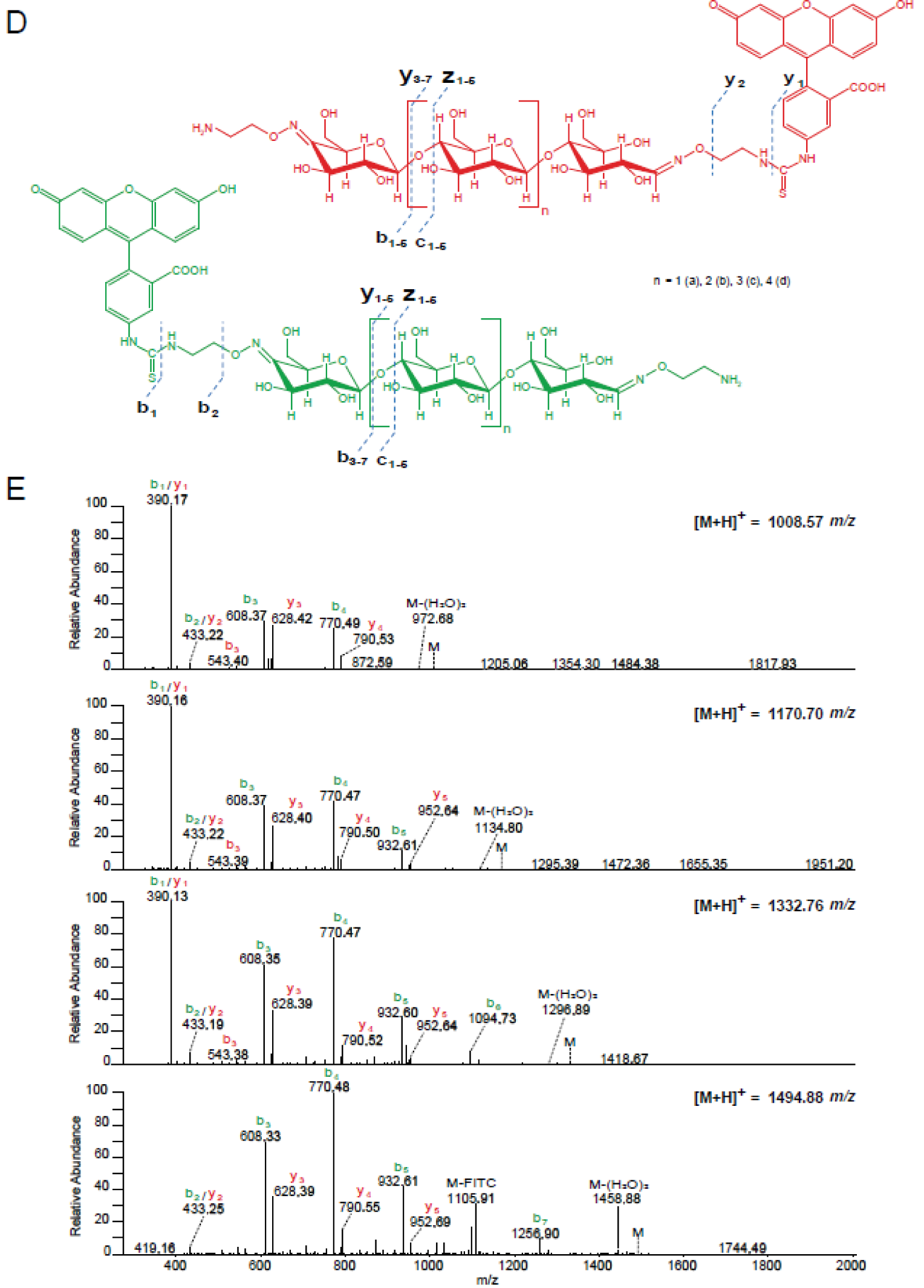
HILIC-FLD-MS analysis of the results of FITC labelling. A) Overlay of extracted ion chromatograms corresponding to single (blue) and double (red) labelled FITC oligosaccharides for *m/z*-ratios a (n=1) 1008.57, b (n=2) 1170.70, c (n=3) 1332.76 and d (n=4) 1494.88 and e (n=1) 1397.87, f (n=2) 1560.00, g (n=3) 1722.09 and h (n=4) 1884.24 respectively. B) MS/MS spectra of the double labelled FITC oligosaccharides. C) Structures of the four double FITC labelled products, with theoretical masses and the observed fragmentation patterns (using the nomenclature of Domon and Costello)^46^. D) Structures of the possible single FITC labelled products arising from incomplete labelling, with theoretical masses and the observed fragmentation patterns (using the nomenclature of Domon and Costello)^46^, with the label located at the reducing end (red) or the non-reducing end (green). E) MS/MS spectra of the single labelled FITC oligosaccharides.

Potential applications of the C4-specific functionalization demonstrated here, include regiospecific immobilization of C4-linked oligosaccharides to solid surfaces such as microarrays, conjugation to protein carriers for immunization purposes, and C4-specific conjugation to nano- or microparticles, or quantum dots^31^. C4-specific functionalization also provides a potential novel analytical tool for studying whether glycosidases act at the reducing end or non-reducing end of oligosaccharides, a feature that has proven challenging to access and may require multistep approaches to resolve^32^. Generating oligosaccharides that are either blocked in the non-reducing end or the reducing end could, in a straightforward way, aid in providing unequivocal evidence of which end is targeted by a given glycosidase. Furthermore, FITC labeling has been shown useful for studying the uptake of specific carbohydrates by microbes^33^. The labeling strategy presented here provides new tools for such studies and could advance these by enabling studies of the directionality of glycan uptake.

Chemo-enzymatic protocols enable synthesis of glycoconjugates with unprecedented precision and are easier than the multistep procedures that are needed when using conventional carbohydrate chemistry. Much progress has been made in recent years on regioselective modification of unprotected sugars^34^, for example through selective chemical oxidation^35^, allowing increasingly efficient chemical synthesis of oligosaccharide-based glycoconjugates. The current study provides a novel chemo-enzymatic route towards glycoconjugates from oligo- and even polysaccharides, providing an attractive alternative to synthesis starting from monosaccharides^36^. We envisage that technologies such as the one presented in this study will help pave the way towards environmentally friendly valorization of biomass and production of useful biochemicals.

## Materials and Methods

### Materials

Cellobiose, cellotriose, cellotetraose, cellopentaose and cellohexaose were from Megazyme (Bray, Ireland). Sodium cacodylate, manganese (II) chloride tetrahydrate, uridine diphosphate galactose (UDP-Gal), fluorescein isothiocyanate (isomer I) and galactosyl transferase from bovine milk were from Sigma. AlexaFluor488 C5-aminooxyacetamide, and bis(triethylammonium) salt were from Invitrogen (Nærum, Denmark). 2-(aminooxy)-1-ethanaminium dichloride was from ABCR GmbH (Karlsruhe, Germany). 2,5-dihydroxy-benzoic acid was from Bruker Daltonics (Bremen, Germany). The LPMO used in this study, from *Neurospora crassa* (*Nc*LPMO9A), and cellobiose dehydrogenase from *Myrococcum thermophilum* (*Mt*CDH) were produced and purified according to Petrovic *et al*. 2019^24^ and Flitsch *et al.* 2013^37^, respectively. The endocellulase Cel5A from *Hypocrea jecorina* was produced according to Saloheimo *et al.* 1988^38^ and the exocelluase Cel6B from *Thermobifida fusca* was produced according to Vuong and Wilson 2009^39^.

### Methods

#### Enzyme reactions

1. *Nc*LPMO9A (1 µM), ascorbic acid (4 mM), phosphoric acid swollen cellulose (PASC, 1% w/v), TrisHCl (10 mM, pH 8.0), 50 °C, 1000 rpm, 12 h.
2. *Mt*CDH (1 µM), *Nc*LPMO9A (1 µM), ascorbic acid (1 mM), phosphoric acid swollen cellulose (PASC, 1% w/v), 50 °C, 1000 rpm, 12 h. *Mt*CDH was used to form double oxidized products.
3. Cel5A (1 µM, endocellulase from *Hypocrea jecorina*), oxime functionalized oligosaccharides (1 mg/mL), 50 °C, 1000 rpm, 12 h.
4. Cel6B (1 µM, exocellulase from *Thermobifida fusca*), oxime functionalized oligosaccharides (1 mg/mL) 50 °C, 1000 rpm, 12 h.

Enzyme reactions 3) and 4) were used to demonstrate that oxime functionalized oligosaccharides were enzymatically degradable.

#### Production of galactosyl-cello-oligosaccharide standards

Galactosyl-cello-oligosaccharides were synthesized using a modified procedure from literature^40^. Briefly, 1 mg of cellotriose or cellotetraose was dissolved in 500 µL of 40 mM sodium cacodylate buffer pH 6.8 containing 40 mM MnCl_2_, 5 mg of UDP-Gal and 0.5 mg of galactosyl transferase from bovine milk (Sigma-Aldrich). The galactosyl transferase will add one galactose to the non-reducing end of the cello-oligosaccharides, forming a β-(1-4) glycosidic bond^41^. The reaction was left to proceed for 3 days at 37 °C with shaking at 1400 rpm. The reaction mixture was left to cool down to room temperature and was directly applied to a CarboGraph column (Grace, Columbia, USA) for desalting of the oligosaccharides, essentially according to a previously published protocol^42^.

#### Functionalization of C4-oxidized cello-oligosaccharides with a hetero-bi-functional linker and labeling with a fluorophore, forming oxime linked products

5 mL of the supernatant from the reactions with PASC (10 mg/mL) prepared as described in enzyme reaction 1 above were used in further experiments. The oligosaccharides were functionalized by reacting carbonyl groups with 2-(aminooxy)-1-ethanaminium dichloride to form oximes, using a modified procedure from literature^43^. The reaction was conducted in 0.1 M NaOAc buffer, pH 4.9, with ca. 50 molar equivalents of the aminooxy linker based on the amount of oxidized oligosaccharides, over a course of 6 days at 37 °C with shaking at 1400 rpm. The products were then purified using active carbon, essentially according to literature^42^. Oxime-functionalized oligosaccharides were labeled with fluorescein isothiocyanate (FITC, ca. 6 molar equivalents with respect to the amino-group content of the oligosaccharide) by incubation in 10 mM sodium bicarbonate buffer, pH 9.5, for 1 hour, with shaking at 1400 rpm, at room temperature and in the dark. The reaction mixture was then freeze-dried.

#### Reduction of soluble products generated by NcLPMO9A

Reduction of oligosaccharides was conducted by mixing 10 µL of oligosaccharide sample from reaction 1 with 65 µL H_2_O (MilliQ) containing 700 µg NaBH_4_ or NaBD_4_. The reduction reaction was incubated over night at ambient temperature and then quenched by adding 20 µL 25 mM sodium acetate. Chromatographic analysis by HPAEC (see below) was done with the sample as is, whereas a further sample preparation for MALDI-ToF-MS was done as follows. Approximately 10 µL of a 1:1 (w/w) suspension of H_2_O:Supelclean ENVI Carb (Sigma-Aldrich) was packed in a pipette tip containing a C8-disc that was trapping the Supelclean material. The bed was conditioned with 50 µL H_2_O. The sample (5 µL) was applied, rinsed with 50 µL H_2_O and then eluted with 20 µL acetonitrile. The eluate was then analyzed by MALDI ToF-MS.

#### HPLC analysis of oligosaccharides

Oligosaccharides were analyzed using three different chromatographical principles. The first principle, high-performance anion-exchange chromatography (HPAEC) at high pH and coupled with pulsed amperometric detection (PAD), was conducted on an ICS3000 system from Dionex (Sunnyvale, California U.S), using previously described conditions and gradients^27,44^. The system was set up with PAD using disposable electrochemical gold electrodes. Two µL samples were injected on a CarboPac PA1 2×250 mm analytical column equipped with a CarboPac PA1 2×50 mm guard column and columns were kept at 30 °C.

The second principle, porous graphitic carbon (PGC) chromatography, was applied using an Ultimate 3000RS (Dionex) UHPLC system, which was connected in parallel (10:1 split) to an ESA Corona Ultra charged aerosol detector (ESA Inc.,Dionex, Sunnyvale, USA) and a Velos pro LTQ linear ion trap (Thermo Scientific, San Jose, CA USA). The PGC column (Hypercarb, 3 µm, 2.1 × 150 mm) including a guard column (Hypercarb, 3 µm, 2.1 × 10 mm) was operated at 70 °C using previously described conditions and gradients^27^. In some cases, 10 µM formic acid was used instead of ammonium acetate, to improve resolution of the oxime coupled products. Two µL samples were injected.

The third principle, hydrophilic interaction chromatography (HILIC), was applied using an Agilent 1290 Infinity (Agilent Technologies, Santa Clara, CA, USA) UHPLC system, which was connected in parallel (1:10 split) to an Agilent 1260 fluorescence detector – Ex 490 nm, Em 520 nm (Agilent Technologies, Santa Clara, CA, USA) and a Velos pro LTQ linear ion trap (Thermo Scientific, San Jose, CA, USA). The HILIC column (bioZen Glycan, 2.6 µm, 2.1 × 100 mm) including a guard column (SecurityGuard ULTRA with bioZen Glycan cartridge, 2.1 × 2 mm) was operated at 50 °C, running at 0.3 mL/min, and using 50 mM ammonium formate pH 4.4 (eluent A) and 100 % acetonitrile (eluent B). Samples were eluted using the following gradient: initial starting ratio of 85 % B and 15 % A, gradient to 75 % B and 25 % A from 0 to 5 mins, gradient to 60 % B and 40 % A from 5 to 25 mins, gradient to 40 % B and 60 % A from 25 to 28 mins, isocratic from 28 to 32 mins, gradient to 85 % B and 15 % A from 32 to 35 mins. Two µL of the samples were injected.

#### MALDI-ToF analysis of oligosaccharides

Two µL of a 9 mg/mL solution of 2,5-dihydroxybenzoic acid in 30% (v/v) acetonitrile was applied to an MTP 384 target plate ground steel TF (Bruker Daltonics). One µL of the sample was then mixed into the DHB droplet followed by drying under a stream of hot air. The samples were analyzed with an Ultraflex MALDI-ToF/ToF instrument (Bruker Daltonics GmbH, Bremen, Germany) with a Nitrogen 337 nm laser beam. The instrument was operated in positive acquisition mode and controlled by the FlexControl 3.3 software package. The acquisition range used was from *m/z* 200 to 7000. The data were collected from averaging 250 laser shots, with the lowest laser energy necessary to obtain sufficient signal to noise ratios. Peak lists were generated from the MS spectra using Bruker FlexAnalysis software (Version 3.3).

#### Mass spectrometry of oligosaccharides, direct injection MS, PGC-MS and HILIC-FLD-MS

For direct injection, oligosaccharides were analyzed using an LTQ-Velos Pro linear ion trap mass spectrometer (Thermo Scientific, San Jose, CA USA) connected to an Ultimate 3000RS HPLC (Dionex, Sunnyvale, California U.S). The setup was used for direct injection without a column; the pump delivered 200 µL/min of 0.03 µM formic acid in 70% acetonitrile and data was acquired for 24 seconds after injection. For the MS, the capillary voltage was set to 3.5 kV and the scan range was *m/z* 150-2000 using two micro scans. The automatic gain control was set to 10,000 charges and the maximum injection time was 20 milliseconds. For fragmentation of selected precursor ions by MS/MS, the normalized collision energy was set to 37 and three micro scans were used. For PGC-MS, the same MS-parameters were used as for direct injection with the exception that the scan range was *m/z* 250-2000. For HILIC-FLD-MS the instrument was operated in positive mode with an ionization voltage of 3.5 kV, auxiliary and sheath gas settings of 5 and 30 respectively (arbitrary units) and with capillary and source temperatures of 300 °C and 250 °C, respectively. The scan range was set to *m/z* 110–2000 and MS/MS analysis was performed with CID fragmentation with helium as the collision gas. All data were recorded with Xcalibur version 2.2.

#### NMR sample preparation

Samples for NMR analysis were lyophilized, then dissolved in 1000 µL 99.9% D_2_O (Chiron, Trondheim, Norway), frozen and lyophilized. The dried samples were then dissolved in 300 µL 99.9% D_2_O and transferred into a 4 mm Shigemi tube (Shigemi, Allison Park, PA, USA).

#### NMR data acquisition and analysis

All homo- and heteronuclear NMR experiments were carried out on a BRUKER Avance III HD 800 MHz spectrometer (Bruker BioSpin AG, Fällanden, Switzerland) equipped with a 5 mm TXI z-gradient probe. For chemical shift assignment, the following spectra were recorded: 1D ^1^H, 2D double quantum filter correlated spectroscopy (DQF-COSY), 2D total correlation spectroscopy (TOCSY) with 70 ms of mixing time, 2D^45^ heteronuclear single quantum coherence (HSQC) with multiplicity editing, 2D^45^ heteronuclear 2-bond correlation (H2BC), 2D [^1^H-^13^C] HSQC-[^1^H,^1^H] TOCSY with 70 ms of mixing time on protons, and 2D [^1^H-^13^C] heteronuclear multiple-bond correlation (HMBC) with BIRD (bilinear rotation decoupling) filtering to suppress first order correlations. These spectra were all recorded at 25 °C. Diffusion-ordered spectroscopy (DOSY) was used to measure the diffusion of the coupled products. A 2D DOSY was set up using a Bruker BioSpin stimulated echo pulse sequence with bipolar gradients (STEBPGP). Gradient pulses of 2 ms duration (δ) and 32 different strengths varying linearly from 0.03 to 0.57 T·m^−1^ were applied and the diffusion delay (Δ) was set to 80 ms. The DOSY spectrum was recorded at 25 °C. The spectra were recorded, processed and analyzed using TopSpin 3.2 software (Bruker BioSpin AG).

## Notes

The authors declare no competing financial interests.

## ACKNOWLEDGEMENTS

This work was supported by the Norwegian Research council through grants 244259, 221576, 226244 and 214613, by the Innovation Fund Denmark, Case No: 0603-00522B, and by the Villum VKR Planet project Nr. 00009283.

---

**Scheme 1.**
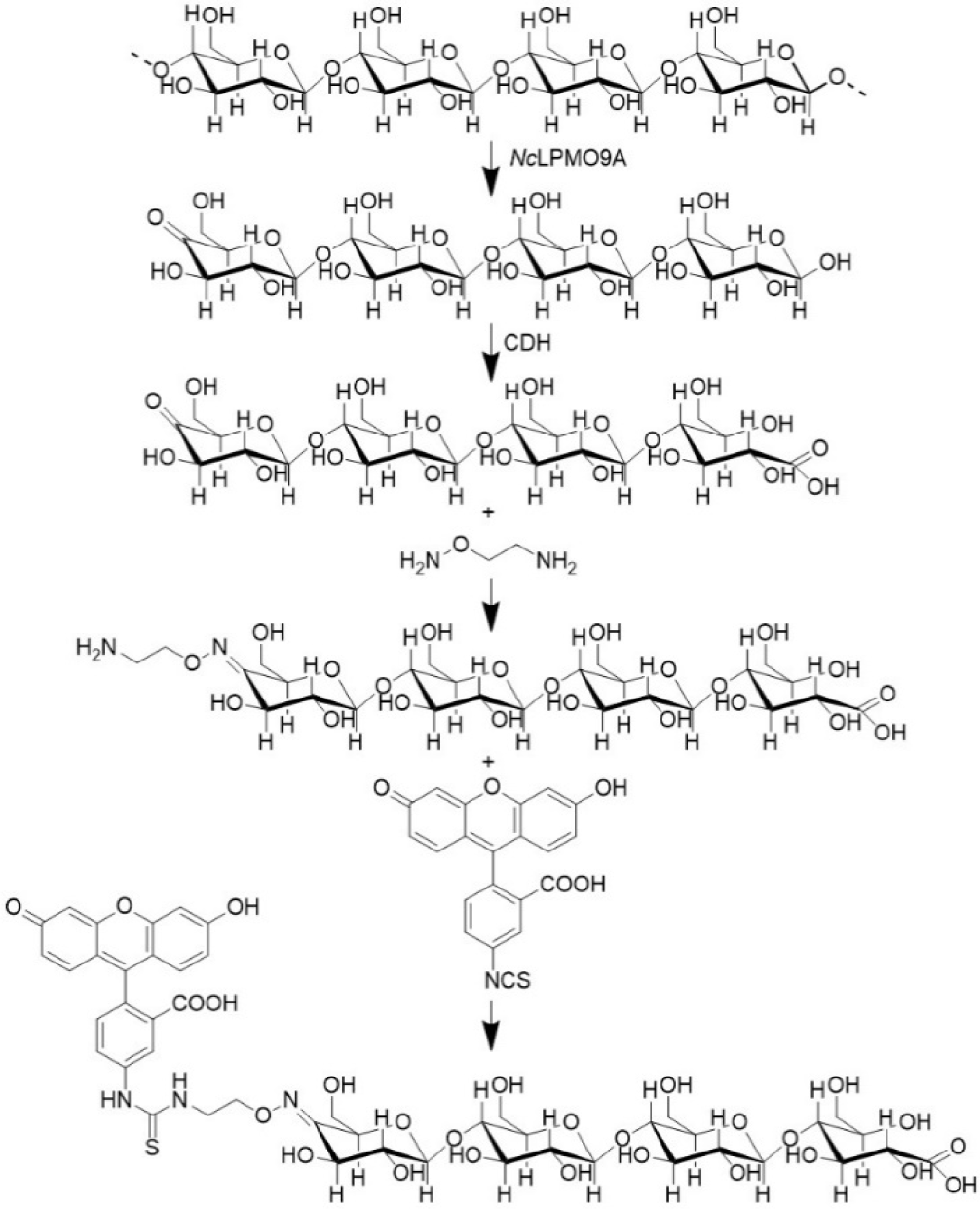
A simple four step chemo-enzymatic procedure to generate a cello-oligosaccharide with FITC conjugated to C4 in the non-reducing end. The starting point is cellulose. *Nc*LPMO9A cuts the cellulose chain and generates C4-oxidized cello-oligosaccharides. CDH converts the reducing end to its corresponding aldonic acid, which protects the reducing end from oxime formation. The oxime linker is coupled to the C-4 keto group prior to the last step in which FITC is cross-linked to the oxime linker.

