## Supplementary material for "Synthesis of Glycoconjugates Utilizing the Regioselectivity of a Lytic Polysaccharide Monooxygenase": Supp

Supplementary Figures

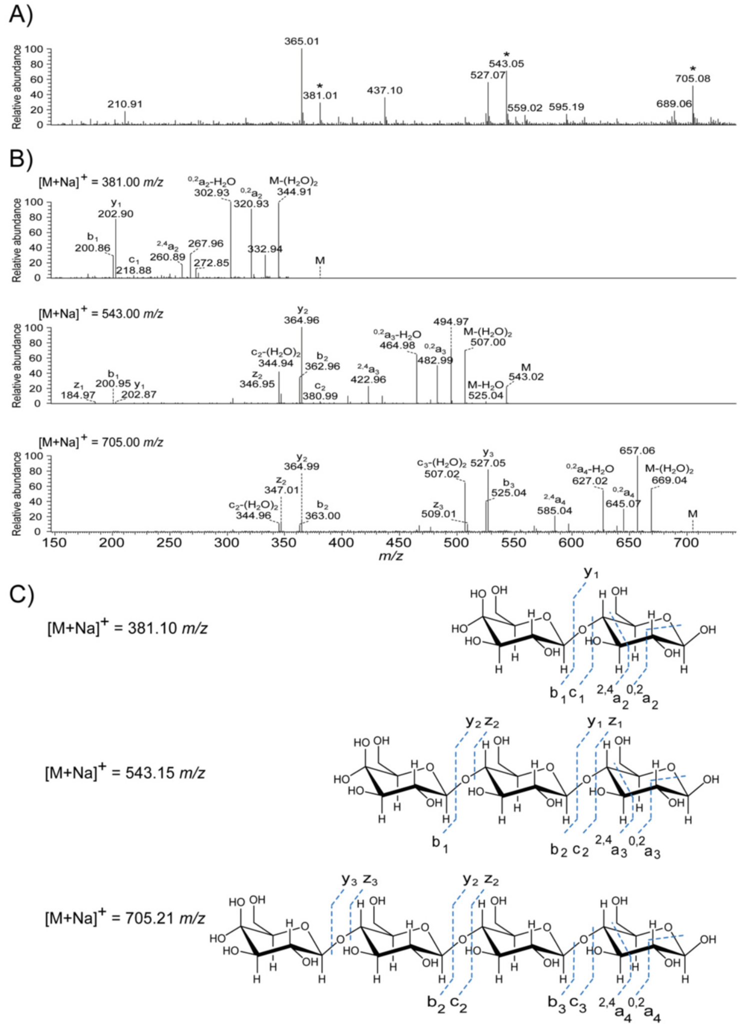


Figure S1. **Direct injection MS analysis of C4-oxidized cello-oligosaccharides.** Oxidized soluble products were analyzed using an LTQ Velos Pro mass spectrometer and direct injection. A) A survey scan of all ions detected in the *m/z*-range 150-750. Peaks annotated with an asterisk are sodium adducts of the hydrated forms of oxidized products and were selected for fragmentation. Note that the keto-sugars are in equilibrium with their gemdiols and that under the conditions used in this study, the keto-form is predominant in MALDI ToF MS analysis (Fig. 1), whereas the gemdiol is predominant in ESI-MS. Peaks corresponding to native cello-oligosaccharides are also visible (*m/z* 365, 527 and 689), which is due to the samples being taken at an early time point in the reaction when there is some release of non-modified sugars from chain ends, while the oxidized oligosaccharides are mostly still bound to the polymer part of the cellulose. B) MS/MS spectra of the three C4-oxidized products show a dominant cleavage of the glycosidic bonds and ring cleavage of the downstream end-sugar. Fragments including the C4-oxidation (a,b,c fragments) tend to lose water as has been described previously[1]. This characteristic double loss of water from the molecular mass is evident for all three products. The MS fragmentation pattern observed here as well as the elution pattern in HPAEC-PAD chromatography are in accordance with previous observations on NMR validated C4-oxidized cello-oligosaccharides[1]. C) The structures of the three C4-oxidized products; the observed fragmentation patterns and the theoretical masses are indicated.

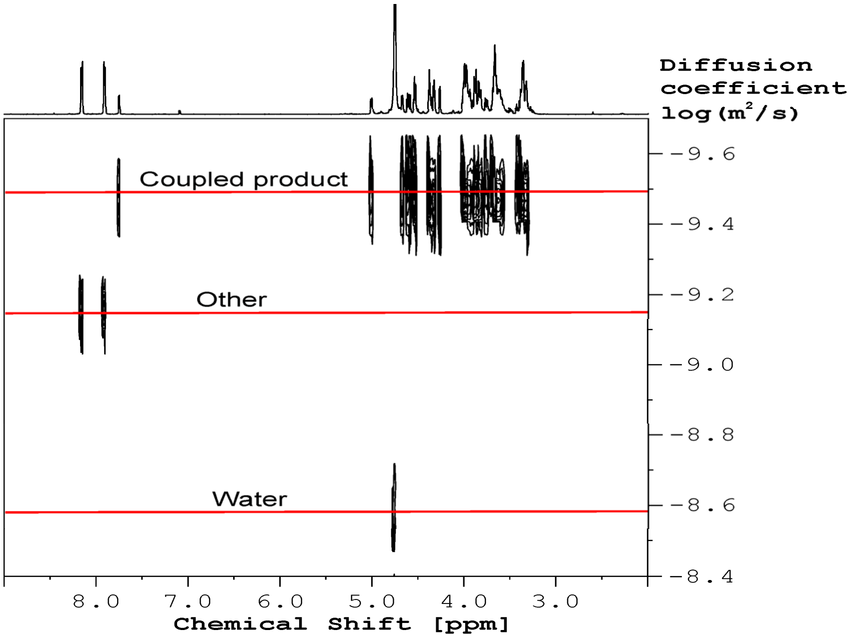


**Figure S2: Diffusion-Ordered Spectroscopy (DOSY) on the aminooxy-linked product (notion coupled product in spectrum)**. DOSY spectrum of the aminooxy-linked product in 99.9 % $D_2O$ recorded at 25°C. Red lines indicate the diffusion of the different molecules in the sample. If the aminooxy-linked products have the same diffusion coefficient it indicates that the C4-oxidized Glc and the oxime linker are covalently linked, while, if they have a different diffusion coefficient, they are not covalently bonded at all. Thus DOSY provides an indirect qualitative method to validate if a coupling reaction has been achieved and helps to identify signals belonging to the aminooxy-linked product (more specifically the azomethine (-C=N-) group which is formed upon coupling). A complete structural elucidation by NMR (Fig.6) was further performed to obtain direct proof of the structure for the coupled product. The [$^1H$-$^{13}C$] HMBC spectrum provided long range bond correlations allowing to connect the monosaccharides for the aminooxy-linked product as well as identification of the carbon chemical shift for a C4-oxidized end at C4 (see figure 6A).

**Table S1: The chemical shift values and correlations of the aminooxy-linked product**. Chemical shifts were assigned for the main product in 99.9% $D_2O$ at 25 °C. Numbers 1-6 represent ring carbon numbers to which the chemical shift values [$^1$H,$^{13}$C] are assigned. The water signal (4.75 ppm) was used as chemical shift reference for $^1$H and $^{13}$C was referenced indirectly using the absolute frequency ratio[2].

| Position | $^1$H [ppm] | $^{13}$C [ppm] | COSY correlations - correlated nuclei | HMBC correlations - correlated nuclei |
|---|---|---|---|---|
| C4ox-1 | 4.67 | 107.0 | C4ox-2 | C4ox-2, C4ox-3, C4ox-5 |
| C4ox-2 | 3.89 | 76.7 | C4ox-1, C4ox-3 | C4ox-1, C4ox-3, C4ox-4 |
| C4ox-3 | 4.27 | 73.4 | C4ox-2 | C4ox-1, C4ox-2, C4ox-4, C4ox-5 |
| C4ox-4 | - | 159.2 | | C4ox-2, C4ox-3, C4ox-5, C4ox-6 |
| C4ox-5 | 5.01 | 77.1 | C4ox-6 | C4ox-1, C4ox-3, C4ox-4, C4ox-6 |
| C4ox-6 | 3.99;3.89 | 63.5 | C4ox-5 | C4ox-5, C4ox-4 |
| Glc3-1 | 4.60 | 104.9 | Glc3-2 | Glc3-2, Glc3-3, Glc3-5 |
| Glc3-2 | 3.39 | 75.6 | Glc3-1, Glc3-3 | Glc3-1, Glc3-3, Glc3-4 |
| Glc3-3 | 3.57 | 77.2 | Glc3-2, Glc3-4 | Glc3-1, Glc3-2, Glc3-4, Glc3-5 |
| Glc3-4 | 3.63 | 81.0 | Glc3-3, Glc3-5 | C4ox-1, Glc3-2, Glc3-3, Glc3-5, Glc3-6 |
| Glc3-5 | 3.64 | 76.7 | Glc3-4, Glc3-6 | Glc3-1,Glc3-4, Glc3-3, Glc3-6 |
| Glc3-6 | 3.95;3.87 | 62.5 | Glc3-5 | Glc3-5, Glc3-4 |
| Glc2-1 | 4.54 | 105.1 | Glc2-2 | Glc2-2, Glc2-3, Glc2-5 |
| Glc2-2 | 3.37 | 75.7 | Glc2-1, Glc2-3 | Glc2-1, Glc2-3, Glc2-4 |
| Glc2-3 | 3.67 | 76.6 | Glc2-2, Glc2-4 | Glc2-1, Glc2-2, Glc2-4, Glc2-5 |
| Glc2-4 | 3.66 | 81.0 | Glc2-3, Glc2-5 | Glc3-1, Glc2-2, Glc2-3, Glc2-5, Glc2-6 |
| Glc2-5 | 3.62 | 77.5 | Glc2-4, Glc2-6 | Glc2-1, Glc2-4, Glc2-3, Glc2-6 |
| Glc2-6 | 3.98;3.84 | 62.4 | Glc2-5 | Glc2-5, Glc2-4 |
| Glc1-1 | 7.76 | 155.4 | Glc1-2 | B', Glc1-2, Glc1-3, Glc1-5 |
| Glc1-2 | 4.61 | 71.9 | Glc1-1,Glc1-3 | Glc1-1, Glc1-3, Glc1-4 |
| Glc1-3 | 3.99 | 73.7 | Glc1-2,Glc1-4 | Glc1-1, Glc1-2, Glc1-4, Glc1-5 |
| Glc1-4 | 3.95 | 80.7 | Glc1-3,Glc1-5 | Glc2-1, Glc1-2,Glc1-3, Glc1-5, Glc1-6 |
| Glc1-5 | 3.96 | 73.6 | Glc1-4,Glc1-6 | Glc1-1, Glc1-3,Glc1-4, Glc1-6 |
| Glc1-6 | 3.88;3.76 | 64.6 | Glc1-5 | Glc1-5 |
| A ($NH_2$-$\mathbf{CH_2}$-$CH_2$-O-R) | 3.34 | 41.4 | B | B |
| A' ($NH_2$-$\mathbf{CH_2}$-$CH_2$-O-R') | 3.29 | 41.2 | B' | B' |
| B ($NH_2$-$CH_2$-$\mathbf{CH_2}$-O-R) | 4.37 | 72.9 | A | B |
| B' ($NH_2$-$CH_2$-$\mathbf{CH_2}$-O-R') | 4.32;4.01 | 72.5 | A' | Glc1-1, A' |
